# Unique viruses that infect Archaea related to eukaryotes

**DOI:** 10.1101/2021.07.29.454249

**Authors:** Ian M. Rambo, Valerie de Anda, Marguerite V. Langwig, Brett J. Baker

## Abstract

Asgard archaea are newly described microbes that are related to eukaryotes. Asgards are diverse and globally distributed, however, their viruses have not been described. Here we characterize seven viral genomes that infected Lokiarchaeota, Helarchaeota, and Thorarchaeota in deep-sea hydrothermal sediments. These viruses code for structural proteins similar to those in *Caudovirales*, as well as proteins distinct from those described in archaeal viruses. They also have genes common in eukaryotic nucleocytoplasmic large DNA viruses (NCLDVs), and are predicted to be capable of semi-autonomous genome replication, repair, epigenetic modifications, and transcriptional regulation. Moreover, Helarchaeota viruses may hijack host ubiquitin systems similar to eukaryotic viruses. This first glimpse of Asgard viruses reveals they have features of both prokaryotic and eukaryotic viruses, and provides insights into their roles in the ecology and evolution of these globally distributed microbes.

---

Asgard archaea are globally distributed microbes that are related to eukaryotes (*1, 2*). Their genomic composition indicates they are descendants of the archaeal host that gave rise to the first eukaryotic common ancestor (*3*). Asgard biodiversity has expanded greatly in recent years due to the recovery of genomes from a range of marine and terrestrial aquatic sediments (*4–6*). An anaerobic, slow-growing Asgard lineage, Lokiarchaeota, has recently been cultured and appears to have syntrophic dependencies with bacteria (*7*). This supports multiple genome-based metabolic inferences of Asgard-bacteria interactions which are thought to have led to the formation of the first mitochondria-containing eukaryotic cell (*7, 8*). It is also hypothesized that interactions with viruses contributed to the origin of complex cellular life. This is based on the presence of nucleus-like viral factories of some NCLDVs (*9*) and bacteriophages (*10, 11*), which allow for replication within the host cytoplasm and the decoupling of transcription and translation by mRNA capping (*12*). Putative viral proteins have been identified within Lokiarchaeota genomes (*13*), suggesting a role of viruses in the exchange of genetic elements and the evolution of Asgards. Type I and Type III CRISPR-Cas immune systems have been described in several Asgard phyla (Odinarchaeota, Thorarchaeota, Lokiarchaeota, and Helarchaeota) (*14*), yet no Asgard-linked virus genomes have been characterized to date.

To explore the role of viruses in Asgard archaea, we searched ~5 TB of assembled metagenomic sequence data from hydrothermal vent-associated sediments in Guaymas Basin (~2,000 m water depth, Gulf of California) (*15, 16*). From this, we recovered 6,756 uncultivated virus genomes (UViGs) (Fig. 1) estimated to be of high- and medium-quality (see Methods). The UViGs were linked to CRISPR spacer sequences from Guaymas Basin metagenome-assembled genomes (MAGs) to determine which viruses infected Asgard cells (*17*). This revealed seven double-stranded DNA (dsDNA) UViGs that infected Lokiarchaeota, Helarchaeota, and Thorarchaeota (Fig. 2, data S1). Interestingly, two Asgard-linked viruses also infected Chloroflexi and Omnitrophica bacteria reconstructed from Guaymas Basin, suggesting a close association of Asgards with these bacteria. One UViG is an integrated provirus in the Chloroflexi genome with CRISPR spacer links to Thorarchaeota, while the other six are non-integrated and classified as lytic. We tentatively name these UViGs after Norse mythological creatures: “Vedfolnir'' (Chloroflexi provirus linked to Thorarchaeota), “Fenrir” (Lokiarchaeota), “Sköll” (Lokiarchaeota), “Nidhogg” (Helarchaeota), and “Ratatoskr” (Helarchaeota and Omnitrophica). Their genome sizes (~21.9-117.5 kilobases) fall within the known range of archaeal dsDNA viruses (*18*) (Fig. 2, data S1). Nidhogg and Fenrir viruses recovered from sample Meg22_1012 have high nucleotide identity to their Meg22_1214 counterparts (99.98 and 99.90% identity, respectively). Nidhogg viruses were closed into complete circular genomes (Fig. 2D; fig. S1; data S2) and the two linear Fenrir viruses are predicted to be >90% complete (data S2).

**Fig. 1.**
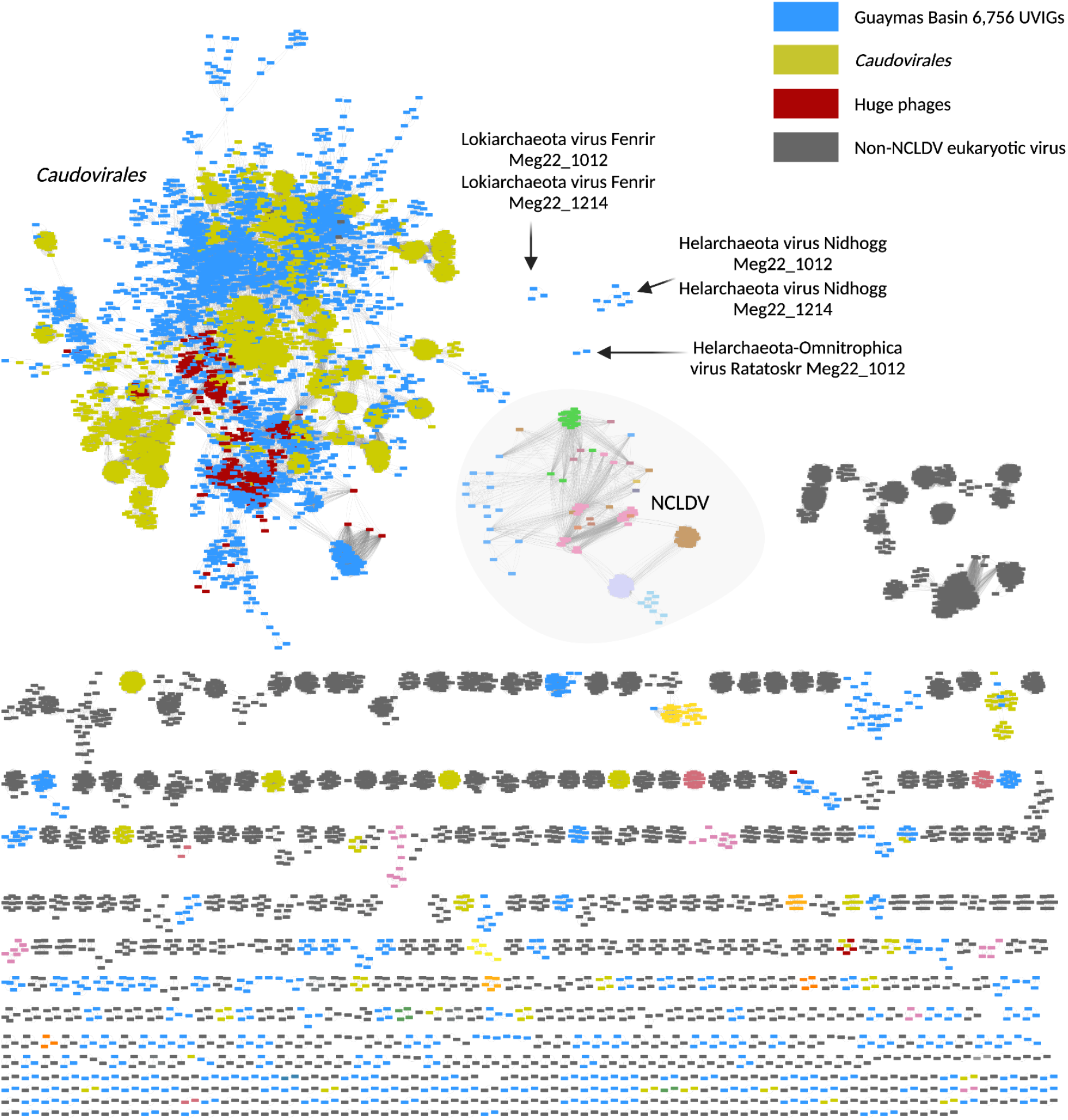
Protein clustering network of virus genomes recovered from Guaymas Basin sediments compared to previously described viruses. A monopartite network analysis constructed with vContact2 v.0.9.19 and Cytoscape v.3.8.0 for taxonomic assignment of Guaymas Basin UViGs (n = 6,756, shown in light blue) was performed using reference eukaryotic, bacterial, and archaeal virus genomes from RefSeq (n = 11,082) and huge phages (n = 361) (see Methods). Nodes represent individual genomes and edges indicate similarity among genomes within a Viral Cluster (n = 770). Asgard-linked viruses are labeled Fenrir, Nidhogg, and Ratatoskr. Vedfolnir and Sköll were classified as outliers.

**Fig. 2.**
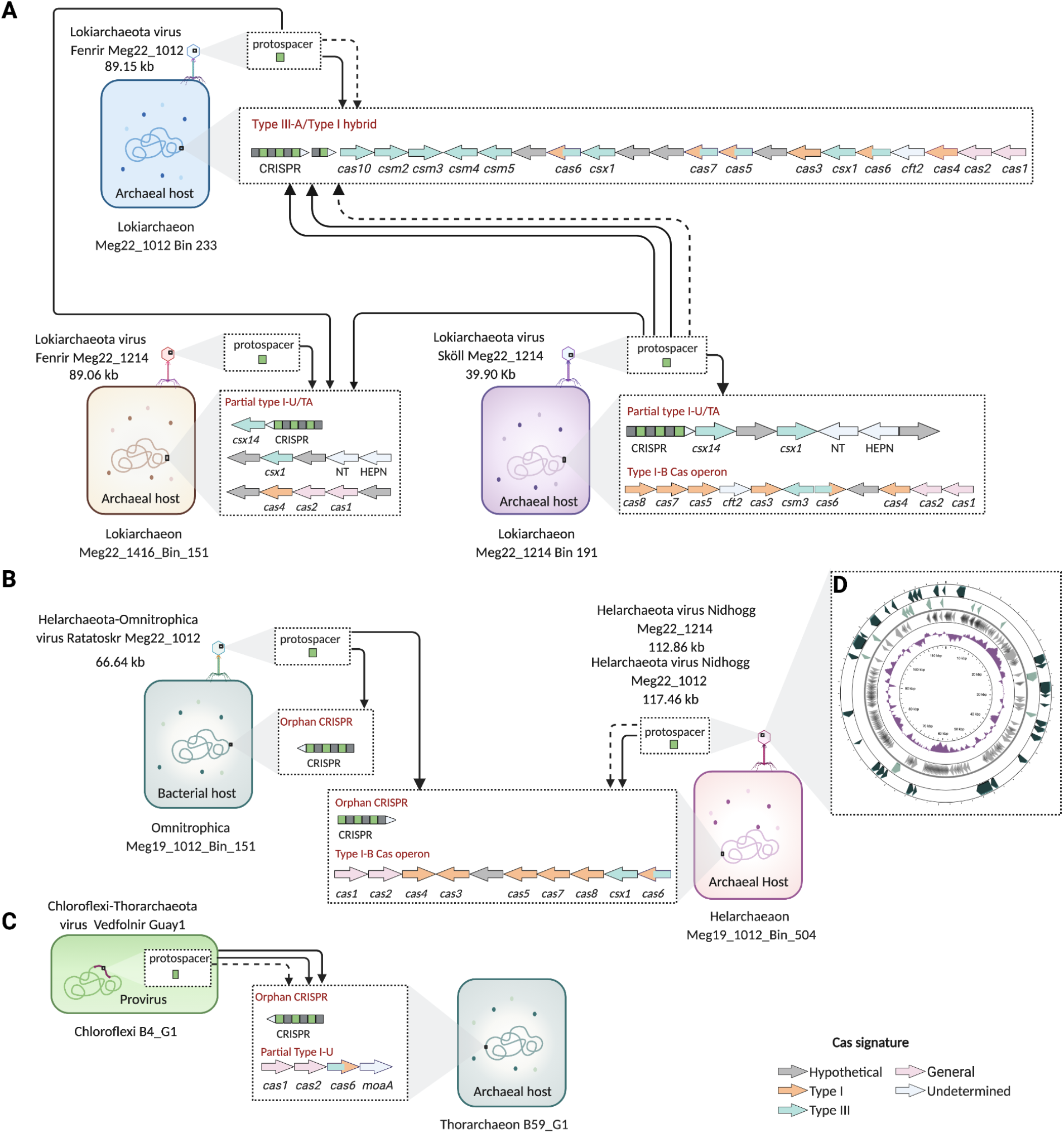
Virus-host linkages to Asgard archaea. (**A**) Lokiarchaeota, (**B**) Helarchaeota, and **(C)**Thorarchaeota MAGs are linked by CRISPR-Cas systems to UViGs. **(D)** Overview of complete circular Nidhogg virus genome. CRISPR spacer blastn-short alignments to a UViG are shown with arrows representing 100% identity (solid) and 95%-99.9% identity (dashed). The Lokiarchaeaota, Helarchaeota, and Omnitrophica MAGs were recovered in this study. The Chloroflexi and Thorarchaeota genomes were recovered previously (*15*). Predicted CRISPR-Cas systems of Asgard MAGs are represented by arrows colored according to the Type signature of their *cas* genes, which point right for (+) and left for (−) sense. Arrows at the end of CRISPR arrays indicate sense in the same fashion. Each *cas* cassette displayed is located on a different scaffold. Viruses are indicated by phage-like icons, although only Ratatoskr contains an identifiable icosahedral capsid. Coding regions within the Nidhogg genome point right for (+) and left for (−) sense, and are drawn to scale. *cas1,* CRISPR-associated endonuclease Cas1; *cas2,* CRISPR-associated endoribonuclease Cas2*; cas3*, CRISPR-associated endonuclease/helicase Cas3*; cas4*, CRISPR-associated exonuclease Cas4*; cas5*, CRISPR system Cascade subunit Cas5; *cas6,* CRISPR-associated endonuclease Cas6*; cas7*, CRISPR-Cas Type I effector complex subunit Cas7*; cas8*, CRISPR-associated protein Cas8*; csm2*, Type III CSM-effector complex small subunit Csm2*; csm3*, Type III RAMP superfamily CSM-effector complex Csm3*; csm4*, Type III RAMP superfamily CSM-effector complex Csm4*; csm5*, Type III RAMP superfamily CSM-effector complex Csm5*; csx1*, CRISPR system endoribonuclease Csx1; *csx14*, Subtype III-U associated protein Csx14; *cft2*, Cft2 family RNA processing exonuclease; *moaA*, molybdenum cofactor biosynthesis protein MoaA; TA, toxin-antitoxin; NT, nucleotidyl transferase; HEPN, higher eukaryotes and prokaryotes nucleotide-binding domain; kb, kilobase. Created with BioRender.com.

In order to classify and understand the evolutionary histories of Asgard viruses, we identified DNA polymerase B (PolB) proteins in the seven UViGs. PolB protein sequences from Fenrir, Nidhogg, and Ratatoskr were aligned with PolB from cellular organisms, phages, archaeal and eukaryotic viruses, and NCLDVs (*19*) (data S3). Phylogenetic analyses of these sequences revealed that Fenrir, Nidhogg, and Ratatoskr are distinct from previously characterized viruses and have a complex evolutionary history (Fig. 3A). The close relationship of Fenrir PolB with Gammaproteobacteria suggests that these genes were horizontally transferred from bacteria, while Nidhogg PolB appears to be phylogenetically related to haloarchaeal myoviruses (Fig. 3A). Ratatoskr PolB is phylogenetically related to Heimdallarchaeota, the Asgard phylum thought to be most closely related to eukaryotes (*2*) (Fig. 3B). To determine the family-level taxonomy of Asgard viruses, we characterized their viral protein family (VPF) composition (*20*) (data S4). All seven Asgard UViGs contain gene orthologs with family-level relatedness to *Myoviridae, Siphoviridae,* and *Podoviridae*, as well as proteins associated with NCLDVs and hyperthermophilic archaeal viruses (Fig. 3C). Asgard UViGs do not cluster with existing reference viruses based on monopartite gene sharing networks (Fig. 1), which may indicate they are novel families or genera of *Caudovirales*. This is supported by previous studies that have identified associations between cosmopolitan *Caudovirales* and archaea (*18, 21*). The lack of clear affiliation with previously described viruses highlights the novelty of these newly recovered Asgard viruses.

**Fig. 3.**
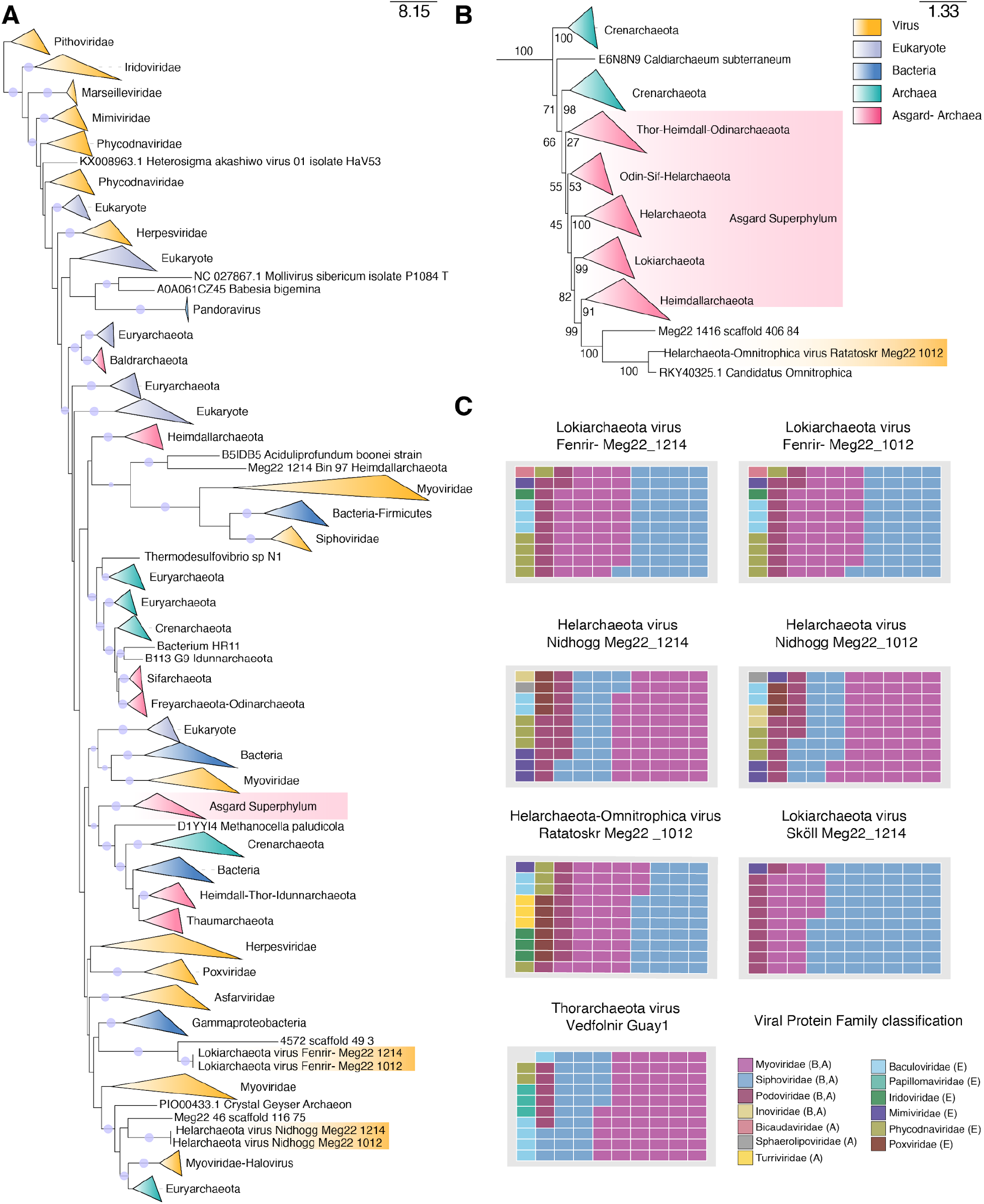
Taxonomic placement of Asgard viruses based on phylogeny and protein composition. (**A**) Maximum-likelihood phylogeny of DNA polymerase B generated using LG+F+R10 model. Circles on tree branches represent ultrafast bootstrap supports >= 95. Viruses described in this paper are highlighted in gold. (**B**) Ratatoskr placement within the LG+F+R10 model tree shown in part A. Bootstraps are shown by values on branches. (**C**) Membership ratios of VPFs shared among Asgard viruses. A, B or E in the Viral Protein Family classification legend indicate archaeal, bacterial, and/or eukaryotic hosts.

We identified hallmark proteins of *Caudovirales* in Asgard UViGs, including minor head protein, baseplate J, tail fibers, portal protein, and terminase large subunit (Fig. 4). Putative capsid proteins which encode a viral protein shell were identified in Ratatoskr, Nidhogg, and Vedfolnir. An HK-97-fold major capsid protein (MCP) was identified in Ratatoskr. Putative MCPs detected in Nidhogg and Vedfolnir (PhANNs, 80% and 90% confidence, respectively) have structural similarities to contractile phage tail sheath-like proteins (>99.9% probability, e-value < 4.4e-24) and phage procapsid protease (99.5% probability, e-value 1.6e-13) (HHPred, data S5). Fenrir and Ratatoskr both encode HNH endonucleases that potentially cleave DNA into genome-length units during packaging, and may operate in concert with their terminase large subunit and portal proteins (*22*). However, putative MCPs were not identified in Fenrir and Sköll genomes, which may be due to the prevalence of novel virion architectures in archaea-specific viruses (*18*). MCPs could be absent due to the tight evolutionary connections of capsid-containing viruses and capsidless selfish elements (CSE). Capsid-containing archaeal viruses have undergone multiple transitions to and from CSEs via the gain and loss of capsid genes as seen in eukaryotic viruses, but still parasitize the genetic information of their hosts (*23, 24*).

**Fig. 4.**
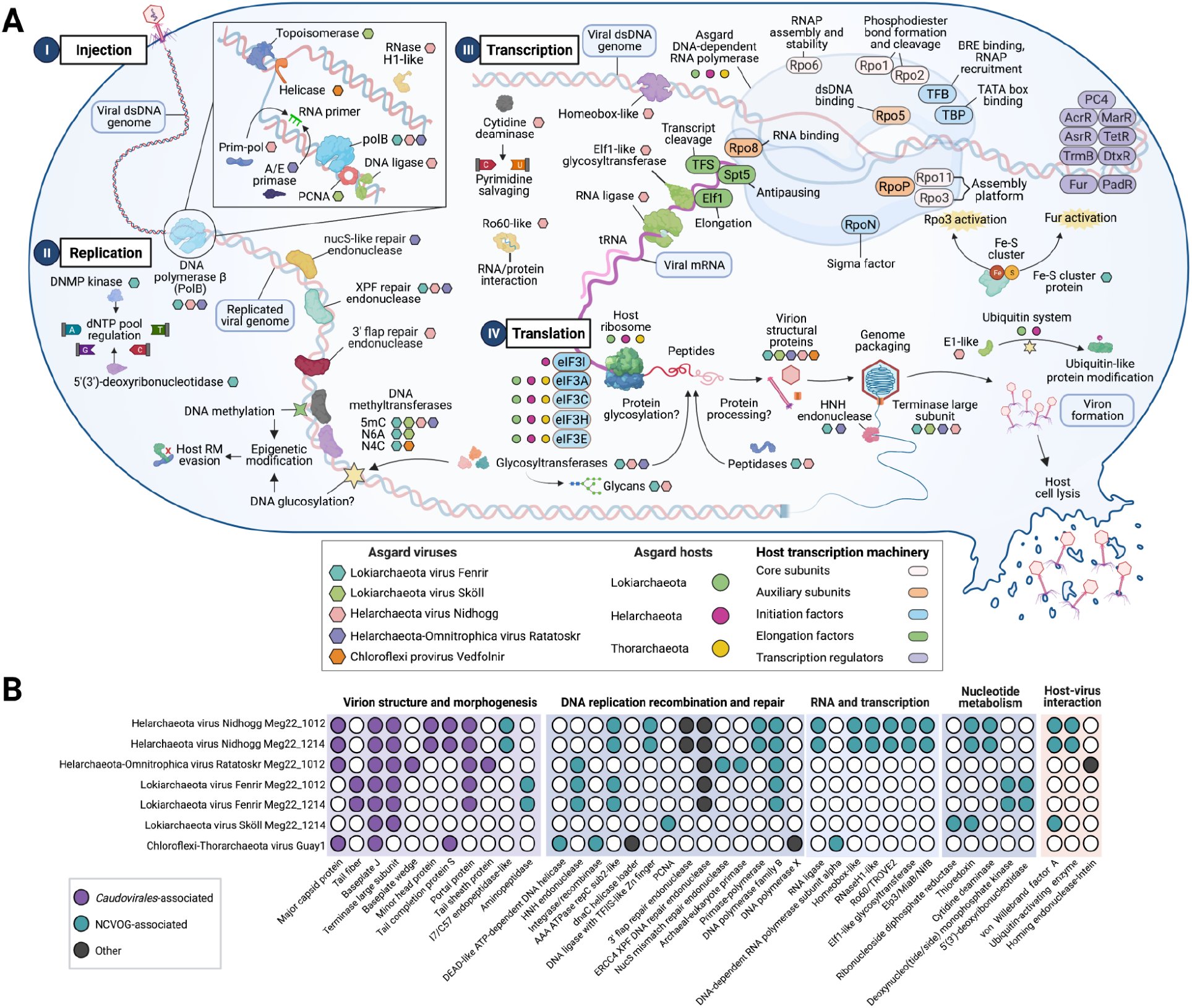
Model of Asgard viral infection mechanisms and protein similarity with other viruses. **(A)** Genes identified in Asgard viruses involved in infection, replication, transcription, and translation. Asgard viruses are shown as different colored hexagons, Asgard hosts as different colored circles, and host transcription machinery is labeled with ovals color coded by function (see legend). **(B)** Distribution of 40 genes (lower x-axis) among Asgard viruses (y-axis). Filled circles indicate annotated orthologs, while white circles indicate that an ortholog was not identified. Abbreviations: NCVOG, Nucleo-Cytoplasmic Virus Orthologous Group (*25*). Created with BioRender.com.

To understand how Asgard viruses impact their hosts, we searched for genes present in prokaryotic viruses and NCLDVs involved in genome replication, nucleotide metabolism, transcription, and host-virus interactions (*19, 25, 26*) (Fig. 4). Asgard viruses encode core DNA replication genes present in prokaryotic and eukaryotic viruses (PolB, archaeo-eukaryotic primase, clamp loaders, RNase H, and ATP-dependent DNA ligases) (*26*). Nidhogg viruses have genes predicted to be involved in autonomous genome replication and proofreading including an ATP-dependent DNA ligase (*ligD*) with a transcription factor S-II-like zinc finger domain, RNase HI/reverse transcriptase-like protein, PolB, and primase-polymerase (PrimPol). Viral PrimPol-like proteins are predicted to have additional roles in DNA and/or RNA priming, as well as damage-tolerant DNA polymerase activity (*27*). These genes have not been described in archaeal viruses to date, and have only been identified in corynebacterium virus BFK20 (*28*) and *Acanthamoeba polyphaga* mimivirus (*29*). The Ratatoskr UViG encodes an archaeal-eukaryotic DNA primase for priming activity in DNA replication (*27*), as well as PolB. Fenrir and Nidhogg encode AAA ATPases with similarity to replication factor C small subunit (*rfcS*) and large subunit (*rfcL*) that may function as clamp loaders. Sköll has proliferating cell nuclear antigen (PCNA) that is most closely related to Lokiarchaeota from deep-sea sediments (36.5% protein similarity). PCNA is an essential replication processivity factor in eukaryotes and archaea (*30*) and is present in some of their viruses (*26*). Sköll also codes for a protein with similarity to a ribonucleotide reductase (ribonucleoside diphosphate reductase, beta subunit) and hedgehog/intein superfamily domains. Ribonucleotide reductase is coded by a conserved core gene for nucleotide metabolism in NCLDVs (*19*). The presence of a hedgehog/intein (Hint) protein domain in this ribonucleotide reductase may indicate that this gene was transferred to Sköll from a previous host. Fenrir viruses may regulate the dNTP pool during replication via a haloacid dehalogenase (HAD) superfamily protein similar to cytosolic 5’(3’) deoxyribonucleotidase. Fenrir also contains a deoxynucleotide monophosphate kinase (DNMP kinase) similar to a large lake algae virus deoxynucleoside monophosphate kinase (bitscore = 115.9, fig. S2) and several *Mimiviridae* DNMP kinases (data S5). Fenrir, Nidhogg, and Ratatoskr viruses possess DNA repair genes, which is a primary characteristic of NCLDVs. This is in contrast to small viruses, which lack these genes (*31*). Fenrir, Nidhogg, and Ratatoskr viruses also encode ERCC4 endonuclease domain proteins with similarity to African swine fever endonuclease EP364R. Nidhogg and Fenrir code ERCC4 endonucleases similar to archaeal XPF 3’ flap repair endonucleases. These XPF-like endonucleases contain helix-hairpin-helix domains, enabling non-specific DNA binding (*32*).

In addition to core DNA replication machinery and repair genes, Fenrir and Nidhogg viruses have a variety of mechanisms for gene expression and post-translational modification. Nidhogg may regulate transcription via a homeodomain protein. Homeodomain proteins are DNA-binding transcription factors present in eukaryotes and *Mimiviridae* (*25, 33*). Nidhogg viruses code a Ro60 protein with TROVE and von Willebrand factor A domains. Ro60 may form a ribonucleoprotein complex with non-coding Y RNA to interact with proteins and other RNAs (*34*). Nidhogg viruses may also repair, splice, and edit transcripts via RNA ligase (*35*) (Fig. 4). Fenrir codes a winged helix-turn-helix domain protein, which is a common DNA binding motif found in archaeal virus transcription factors (*21*). We did not identify any translation machinery in these UViGs, suggesting they utilize host ribosomes and eukaryotic initiation factors present in the Asgard hosts. However, Nidhogg and Fenrir may process viral protein precursors with encoded peptidases. Nidhogg is predicted to code a peptidase with similarity to Vaccinia virus I7L/C57-family processing endopeptidase, while Fenrir appears to be capable of producing an M42-family aminopeptidase (Fig. 4).

Several Asgard phyla code homologs of eukaryotic ubiquitin protein modification systems (*2*). Eukaryotic viruses have been shown to hijack these systems to aid in various stages of viral propagation (*36*). Nidhogg codes three ubiquitin-activating (ThiF family/E1-like) protein homologs (fig. S3) that could harness Helarchaeota ubiquitin systems (E1-like activating enzyme, E2-like conjugating enzyme, and ubiquitin-like protein domain) (data S6) and repurpose them for processes such as replication, transcription, viron formation, and release (*36*). Phylogenetic analyses of Nidhogg proteins suggest they are novel E1 ubiquitin-activating homologs that are ancestral to Helarchaeota E1-like and eukaryotic ubiquitin-like modifier activating 5 proteins (fig. S1; data S7). This is interesting given that this type of host-virus interaction has only been described in eukaryotic systems.

Asgard viruses contain genes for autonomous DNA methylation and glycosylation that may play roles in host interactions. DNA methylation has been documented in bacteriophages (*37*) and *Mimiviridae* (*38*) as an epigenetic modification used to evade host defenses like restriction-modification (RM) systems, hijack transcription machinery, regulate replication cycles, and/or defend against competing viruses with RM systems. Interestingly, the Asgard viruses in Guaymas Basin may be capable of autonomous DNA methylation, as they code for 5-methylcytosine, N4-methylcytosine, and D12 class N6-methyladenine methyltransferases. The concurrence of restriction endonuclease (RE)-like genes, such as a RecB family RE in Nidhogg, may indicate functional RM systems that could inhibit host gene expression and shift transcription from host to virus (*39*). Fenrir, Nidhogg, and Ratatoskr also encode hexosyltransferases, a class of glycosyltransferases (GTs) that may confer the ability to modify hexoses on DNA, proteins, or lipids in order to evade host defense systems and inhibit host molecular functions (*40*). GTs have been identified in phages and NCLDVs (*40, 41*), but are not well-documented in archaeal viruses. Fenrir encodes two overlapping genes with sequence homology to *Sulfolobus islandicus* rod-shaped virus 1 uncharacterized GT (SIRV1-GT) and alpha-1,2-mannosyltransferase (WbdA). This suggests that Fenrir could produce mannose glycan chains that are terminated by an encoded SAM-dependent methyltransferase (WbdD) (*42*) without utilizing host machinery. Additionally, Fenrir encodes L-malate GT (BshA), a gene that is prevalent in Asgards and is thought to be involved in the biosynthesis of low molecular weight thiols (LMWT) in bacteria and archaea (*43*). Ratatoskr encodes an N-acetyl-alpha-D-glucosaminyl L-malate dehydrogenase 1 (BshB1) homolog, a gene that is also implicated in LMWT biosynthesis. Finally, Nidhogg encodes a GT with hits to SIRV1-GT, as well as a glycogen synthase (GlgA) homolog which may form glucan chains (Fig. 4A).

Given the relatedness of Asgards to eukaryotes, it is interesting that Asgard-infecting viruses have replication and regulation machinery akin to eukaryotic NCLDVs (PCNA, PolB, ribonucleotide reductase, DNA ligase, DNMP kinase, and replication factor C-like AAA ATPases). However, mRNA capping genes that allow for uncoupling of transcription and translation in NCLDVs do not appear to be present in these viruses. In addition, many of their characteristics are distinct from previously studied archaeal viruses. For example, the complete Nidhogg genomes code for homeobox and PrimPol protein domains which are lacking in archaeal viruses. They also contain genes for ubiquitin protein modification systems that have not been previously described in archaeal viruses, which are used for viral propagation in eukaryotic viruses. They code structural proteins reminiscent of those in *Caudovirales*, yet most of their protein composition is distinct from this order. Asgard viruses appear to have both archaeal and eukaryotic viral characteristics, which is consistent with the evolutionary position of their hosts. This first description of Asgard-linked viruses advances our understanding of the roles of viruses in the ecology and evolution of Asgards.

## Acknowledgments

This work was supported by the Moore-Simons Project on the Origin of the Eukaryotic Cell, Simons Foundation 73592LPI (https://doi.org/10.46714/735925LPI) and Simons Foundation early career award 687165 to B.J.B. Sampling in Guaymas Basin (Gulf of California) was supported by NSF Awards OCE-0647633 to Dr. Andreas P. Teske. We thank Dr. Daniel Tamarit and Dr. Thijs Ettema for advice about this research.

## Data and materials availability

The genomic sequences associated with the study have been deposited in NCBI under BioProject PRJNA743331.

## Materials and Methods

### Metagenomic assembly and binning

The sampling, assembly, and binning methods for the genomic data analyzed in this study are described in detail in references (*15, 16*). Samples were collected from Guaymas Basin (GB) in the Gulf of California, Mexico (27°N 0.388, 111°W 24.560) from a depth of ~2,000 m using polycarbonate cores. Whole community DNA from ≥10 g of sediment was extracted using the DNeasy PowerSoil and MoBio PowerMax kits. DNA concentrations were quantified using a QUBIT 2.0 fluorometer (Thermo-Fisher, Singapore) and metagenomic sequencing was performed at the Michigan State University RTSF Genomics Core and the Joint Genome Institute. Sequencing was performed in a 2×150 bp and 2×125 bp paired end format. Sequences described in (*16*) were trimmed and quality controlled using Sickle v1.33 (*44*) and assembly was performed using IDBA-UD v1.0.9 (--seed_kmer 55) (*45*). In (*15*), sequences were quality controlled and assembled by JGI. In (*16*) binning of individual GB assemblies (scaffolds >2 000 bp) was performed using Concoct v0.4.0 (*46*) and Metabat v2.12.1 (*47*). Concoct was used with default settings and Metabat was run with the following parameters: –minCVSum 0 --saveCls -d -v --minCV 0.1 -m 2000. Consensus MAGs produced from these two binning tools were determined using DAS Tool v1.0 (*48*) with default settings. In (*15*) binning was performed using ESOM, Anvi’o, and Metabat, and results from all three binning tools were combined using DAS Tool v1.0. CheckM v1.0.11 and v1.0.5 (*49*) was used to determine MAG completeness and contamination (Table S2). Binned and unbinned contigs >= 2 kb were annotated with IMG/M v5.0 (Chen et al 2019). Annotations were consolidated and mapped to MAGs using the MetaGaia metabolic_profile.py script (https://github.com/valdeanda/MetaGaia). Additional annotation of Asgard MAGs was performed with InterProScan v5.31-70.0 (*50*).

### CRISPR-Cas system prediction and subtyping

CRISPR arrays were predicted in Guaymas MAGs with CRISPRDetect v.2.4 (*51*) using the following parameters: -minimum_no_of_repeats 3 -minimum_repeat_length 23 -right_flank_length 500 -left_flank_length 500 -array_quality_score_cutoff 3. Spacer sequences were extracted from the output GFF3 file using a custom Python script, and de-replicated using CD-HIT-EST v4.8.1 (*52*) using the following parameters: -c 1.0 -n 11 -b 20. CRISPRDetect and IMG/M CRISPR Recognition Tool (*53*) results were compared to verify CRISPR array prediction.

Cas gene cassette prediction and subtyping was performed for MAGs using CRISPRcasIdentifier (*54*), with combinations of Classification and Regression Trees (CART), Support Vector Machine (SVM), and Extremely Randomized Trees (ERT) algorithms for classification and regression. Subtypes were assigned via a consensus-based approach in which all combinations of CART, SVM, and ERT classifiers/regressors provided the same subtype assignment, with that subtype probability exhibiting the highest mean and lowest standard deviation of the predictions. *Cas* cassette subtype prediction annotations were compared against IMG/M and InterProScan results, and subtype predictions were verified by ensuring the presence of key effector complex genes.

### Viral contig recovery and annotation

VIBRANT v.1.2.1 was used to recover uncultivated viral genomes (UViGs) as previously described (*55*). In brief, contigs >10 kb with >4 open reading frames (ORFs) predicted using Prodigal v2.6.3 (*56*) were utilized for annotation against KOALA-KOFAM v2019-03-20 (*57*), Pfam v32 (*58*), and virus orthologous groups (VOG) release 94 (*59*) databases with hmmsearch via HMMER v3.3.2 (*60*). For contigs containing a provirus, the proviral region was extracted and retained if it exhibited a clear shift from the host genome, met the aforementioned sequence length and ORF requirements, and contained a putative integrase. Lytic and provirus UViGs designated in VIBRANT as mid- and high quality UViGs based on Virus Orthologous Groups (VOGs), nucleotide replication proteins, and enrichment of viral hallmark genes (e-value and bitscore cutoffs of 1e-5 and 30, respectively) were retained. Asgard virus genome completeness and quality was additionally assessed with CheckV v0.8.1 (*61*) using HMM and amino acid identity (AAI) metrics (end_to_end mode; CheckV database v1.0). Additional gene annotation was performed with DIAMOND v.2.0.4.142 (*62*) using BLASTP (*63*) against the NCBI NR database (downloaded October 4, 2020). HMM-based annotations were completed using InterProScan v5.31-70.0 (*50*), IMG/M v5.0 (*64*), and nucleocytoplasmic virus orthologous group HMMs (NCVOGs) (*65*) with HMMER v3.3.2 (*60*). Artificial neural network-based classifications were performed with the PhANNs v0.3 web server (*66*). Candidate UViGs were verified to contain hypothetical gene enrichments. Protein homology of putative major capsid proteins assigned by PfANNs was assessed using the HHpred web server (*67*) against the PDB_mmCIF70_17_May, SCOPe70_2.07, Pfam-Av34, and UniProt-SwissProt-viral70_23_August_2020 databases (HHBlits=>UniRef30 MSA generation method, 3 MSA generation iterations, 20% min coverage of MSA hits, local alignment, no MAC realignment, secondary structure scoring during alignment, 1e-6 cutoff for MSA generation, 0% min seq identity of MSA hits with query). Visualization of the circular Nidhogg genome was performed with the CGView server (*68*).

### Protein clustering and taxonomic comparisons against publicly available viruses

To cluster GB UViGs for family and genus-level taxonomic assignment against known viruses, protein clustering networks were utilized with vContact2 v.0.9.19 (*69*) using all-vs-all DIAMOND alignments, ClusterONE (*70*) and MCL (*71*) (parameters --pcs-mode MCL --vcs-mode ClusterONE --seed-method unused_nodes --similarity match --merge-method single --min-size 2 --rel-mode Diamond) with the RefSeq v201 prokaryotic virus database, eukaryotic viruses from RefSeq v202 viral database (*72*) and huge phage genomes (*73*). Protein clustering networks were visualized in Cytoscape v3.8.0 (*74*) using an edge-weighted spring embedded model. UViG family and genus-level taxonomic classification were performed with VPF-Class v1.0 (*20*) via reference viral protein families (VPFs) using a minimum e-value cutoff of 1e-4. VPF classification files from May 20, 2020 were used to build the reference database. Waffle charts representing VPF membership ratios were created in RStudio v1.3.959 (*75*) using R v4.0.0 (*76*) and the ‘waffle’ package v1.0.1 (*77*).

### CRISPR-based host-virus linking

Host-virus infections can be resolved by searching for CRISPR proto-spacer sequences in the host that match those present in the viral community. To do this, spacers were aligned to viral contigs using blastn-short withvia BLAST+ v2.5.0 (*63*) (percent identity >= 95%, e-value <= 1e-5, 100% query coverage, word size = 8, maximum target sequences = 10,000,000, allowing 1 mismatch). An undirected network of significant virus-host CRISPR spacer matches was constructed with NetAn v.1.0 (*78*) and visualized in RStudio v.1.3.959 (*75*) with VisNetwork v.2.0.9 (https://github.com/datastorm-open/visNetwork).

### Phylogenetic analysis

A phylogenetic tree of DNA polymerase family B (PF00136) was constructed to better resolve the evolutionary relationships of Asgard-linked viruses. This protein is broadly distributed among all domains of life and DNA viruses. *Caudovirales* and *Herpesvirales* sequences from PF00136 were de-replicated at 100% similarity using CD-HIT (*52*) (word-size of 5). Bacterial sequences were clustered at 50% similarity with CD-HIT (word-size of 3). Eukaryote, NCLDV, and archaea references were selected as previously described (*79*). A total of 746 sequences were aligned with MAFFT v7.457 (*80*). Poorly-aligned sequences were manually removed with Geneious Prime v2021.0.3 and re-aligned with MAFFT v7.450. Alignment columns comprising >50% gaps were removed. Maximum-likelihood phylogenetic trees were constructed with IQTree v2.0.3 (*81*) and model selection was performed with ModelFinderPlus (*82*). Two trees were constructed as follows: 1) LG+F+R10 with 1,000 ultrafast bootstraps (parameters: -m MFP -bb 1000 -bnni -alrt 1000 -wsl -wsr -wbt -mset LG,LG+C20,LG+C40,LG+C60,LG4M,LG4X -mrate,R4,R5,R8,R10,G4,G8 -mfreq ,F -nt 60) and 2) LG+F+R10 with the C60 complex mixture model over 1,000 ultrafast bootstraps (parameters: -m MFP -bb 1000 -bnni -alrt 1000 -wsl -wsr -wbt -mset LG,LG+C20,LG+C40,LG+C60,LG4M,LG4X -mrate,R4,R5,R8,R10,G4,G8 -mfreq ,F -nt 60), followed by a posterior mean site frequency model approximation using the ultrafast bootstrap guide trees and LG+C60+F+R10 model.

A maximum likelihood phylogeny of DNMP kinase was constructed to resolve the placement of Lokiarchaeota virus Fenrir sequences. Reference bacterial and viral DNMP kinase sequences were downloaded from Uniprot, along with top BLASTP hits to Fenrir virus DNMP kinase (word size = 6, 1e-10 maximum e-value, bitscore >= 90). Uniprot sequences were clustered at 97% identity (for bacteria) and de-replicated at 100% identity (for viruses) with CD-HIT (word size = 5). Sequences were aligned with MUSCLE v3.8.1551. The MSA was curated in Geneious Prime v2021.0.3 by manually removing poorly-aligned sequences and re-aligning with MAFFT. Alignment columns comprising >50% gaps were removed. A maximum-likelihood phylogenetic tree of 241 sequences was constructed with RAxML v8.2.11 (HPC-PTHREADS-AVX, parameters: -T 30 -m PROTGAMMAAUTO -N 1000 -p 44 -x 44 -f a) using an LG likelihood model with fixed base frequencies.

The ubiquitin-activating enzyme E1-like tree was constructed from 11 protein sequences belonging to the NEDD8-activating enzyme E1 catalytic subunit family (IPR030468), 218 sequences belonging to the ubiquitin-activating E1 enzyme (IPR035985), 14 viral sequences obtained from NCBI (search parameters: ThiF family protein[Protein Name] OR ubiquitin-activating enzyme [Protein Name] AND viruses[Filter]), 133 sequences derived from Lokiarchaeota and Helarchaeota based on InterProScan v5.31-70.0 results, and 3 Nidhogg virus sequences. Sequences were aligned with MAFFT v7.457 (*80*). The MSA was curated in Geneious Prime v2021.0.3 by manually removing poorly-aligned sequences and re-aligning with MAFFT v7.457. Alignment columns comprising >50% gaps were removed. The phylogeny was constructed with IQTree v1.6.1 (*81*) and the LG+R8 model was selected with ModelFinderPlus (*82*) (parameters: -m MFP -bb 1000 -bnni -seed 42 -nt 40). All phylogenetic trees described in this study were visualized in iTol v6.3 (*83*)

## References

1. A. Spang, A. J. H. Saw, S. L. Jørgensen, K. Zaremba-Niedzwiedzka, J. Martijn, A. E. Lind, R. van Eijk, C. Schleper, L. Guy, T. J. G. Ettema, Complex archaea that bridge the gap between prokaryotes and eukaryotes. Nature. 521, 173–179 (2015).

2. K. Zaremba-Niedzwiedzka, E. F. Caceres, J. H. Saw, D. Bäckström, L. Juzokaite, E. Vancaester, K. W. Seitz, K. Anantharaman, P. Starnawski, K. U. Kjeldsen, M. B. Stott, T. Nunoura, J. F. Banfield, A. Schramm, B. J. Baker, A. Spang, T. J. G. Ettema, Asgard archaea illuminate the origin of eukaryotic cellular complexity. Nature. 541, 353–358 (2017).

3. L. Eme, A. Spang, J. Lombard, C. W. Stairs, T. J. G. Ettema, Archaea and the origin of eukaryotes. Nat. Rev. Microbiol. 15, 711–723 (2017).

4. K. W. Seitz, C. S. Lazar, K.-U. Hinrichs, A. P. Teske, B. J. Baker, Genomic reconstruction of a novel, deeply branched sediment archaeal phylum with pathways for acetogenesis and sulfur reduction. ISME J. 10, 1696–1705 (2016).

5. K. W. Seitz, N. Dombrowski, L. Eme, A. Spang, Asgard archaea capable of anaerobic hydrocarbon cycling. Nature (2019) (available at https://www.nature.com/articles/s41467-019-09364-x).

6. B. J. Baker, V. De Anda, K. W. Seitz, N. Dombrowski, A. E. Santoro, K. G. Lloyd, Diversity, ecology and evolution of Archaea. Nat Microbiol. 5, 887–900 (2020).

7. H. Imachi, M. K. Nobu, N. Nakahara, Y. Morono, M. Ogawara, Y. Takaki, Y. Takano, K. Uematsu, T. Ikuta, M. Ito, Y. Matsui, M. Miyazaki, K. Murata, Y. Saito, S. Sakai, C. Song, E. Tasumi, Y. Yamanaka, T. Yamaguchi, Y. Kamagata, H. Tamaki, K. Takai, Isolation of an archaeon at the prokaryote–eukaryote interface. Nature. 577, 519–525 (2020).

8. A. Spang, C. W. Stairs, N. Dombrowski, L. Eme, J. Lombard, E. F. Caceres, C. Greening, B. J. Baker, T. J. G. Ettema, Proposal of the reverse flow model for the origin of the eukaryotic cell based on comparative analyses of Asgard archaeal metabolism. Nat Microbiol. 4, 1138–1148 (2019).

9. P. Forterre, M. Gaïa, Giant viruses and the origin of modern eukaryotes. Curr. Opin. Microbiol. 31, 44–49 (2016).

10. V. Chaikeeratisak, K. Nguyen, K. Khanna, A. F. Brilot, M. L. Erb, J. K. C. Coker, A. Vavilina, G. L. Newton, R. Buschauer, K. Pogliano, E. Villa, D. A. Agard, J. Pogliano, Assembly of a nucleus-like structure during viral replication in bacteria. Science. 355, 194–197 (2017).

11. L. M. Malone, S. L. Warring, S. A. Jackson, C. Warnecke, P. P. Gardner, L. F. Gumy, P. C. Fineran, A jumbo phage that forms a nucleus-like structure evades CRISPR-Cas DNA targeting but is vulnerable to type III RNA-based immunity. Nat Microbiol. 5, 48–55 (2020).

12. P. J. L. Bell, Evidence supporting a viral origin of the eukaryotic nucleus. Virus Res. 289, 198168 (2020).

13. M. Krupovic, V. V. Dolja, E. V. Koonin, The LUCA and its complex virome. Nat. Rev. Microbiol. 18, 661–670 (2020).

14. K. S. Makarova, Y. I. Wolf, J. Iranzo, S. A. Shmakov, O. S. Alkhnbashi, S. J. J. Brouns, E. Charpentier, D. Cheng, D. H. Haft, P. Horvath, S. Moineau, F. J. M. Mojica, D. Scott, S. A. Shah, V. Siksnys, M. P. Terns, Č. Venclovas, M. F. White, A. F. Yakunin, W. Yan, F. Zhang, R. A. Garrett, R. Backofen, J. van der Oost, R. Barrangou, E. V. Koonin, Evolutionary classification of CRISPR-Cas systems: a burst of class 2 and derived variants. Nat. Rev. Microbiol. 18, 67–83 (2020).

15. N. Dombrowski, A. P. Teske, B. J. Baker, Expansive microbial metabolic versatility and biodiversity in dynamic Guaymas Basin hydrothermal sediments. Nat. Commun. 9, 4999 (2018).

16. C. J. Castelle, R. Méheust, A. L. Jaffe, K. Seitz, X. Gong, B. J. Baker, J. F. Banfield, Protein Family Content Uncovers Lineage Relationships and Bacterial Pathway Maintenance Mechanisms in DPANN Archaea. Front. Microbiol. 12, 660052 (2021).

17. A. F. Andersson, J. F. Banfield, Virus population dynamics and acquired virus resistance in natural microbial communities. Science. 320, 1047–1050 (2008).

18. D. Prangishvili, D. H. Bamford, P. Forterre, J. Iranzo, E. V. Koonin, M. Krupovic, The enigmatic archaeal virosphere. Nat. Rev. Microbiol. 15, 724–739 (2017).

19. L. M. Iyer, L. Aravind, E. V. Koonin, Common origin of four diverse families of large eukaryotic DNA viruses. J. Virol. 75, 11720–11734 (2001).

20. J. C. Pons, D. Paez-Espino, G. Riera, N. Ivanova, N. C. Kyrpides, M. Llabrés, VPF-Class: Taxonomic assignment and host prediction of uncultivated viruses based on viral protein families. Bioinformatics (2021), doi:10.1093/bioinformatics/btab026.

21. M. Krupovic, V. Cvirkaite-Krupovic, J. Iranzo, D. Prangishvili, E. V. Koonin, Viruses of archaea: Structural, functional, environmental and evolutionary genomics. Virus Res. 244, 181–193 (2018).

22. S. Kala, N. Cumby, P. D. Sadowski, B. Z. Hyder, V. Kanelis, A. R. Davidson, K. L. Maxwell, HNH proteins are a widespread component of phage DNA packaging machines. Proc. Natl. Acad. Sci. U. S. A. 111, 6022–6027 (2014).

23. J. Iranzo, P. Puigbò, A. E. Lobkovsky, Y. I. Wolf, E. V. Koonin, Inevitability of Genetic Parasites. Genome Biol. Evol. 8, 2856–2869 (2016).

24. Koonin Eugene V., Dolja Valerian V., Virus World as an Evolutionary Network of Viruses and Capsidless Selfish Elements. Microbiol. Mol. Biol. Rev. 78, 278–303 (2014).

25. N. Yutin, Y. I. Wolf, D. Raoult, E. V. Koonin, Eukaryotic large nucleo-cytoplasmic DNA viruses: clusters of orthologous genes and reconstruction of viral genome evolution. Virol. J. 6, 223 (2009).

26. D. Kazlauskas, M. Krupovic, Č. Venclovas, The logic of DNA replication in double-stranded DNA viruses: insights from global analysis of viral genomes. Nucleic Acids Res. 44, 4551–4564 (2016).

27. T. A. Guilliam, B. A. Keen, N. C. Brissett, A. J. Doherty, Primase-polymerases are a functionally diverse superfamily of replication and repair enzymes. Nucleic Acids Res. 43, 6651–6664 (2015).

28. N. Halgasova, I. Mesarosova, G. Bukovska, Identification of a bifunctional primase-polymerase domain of corynephage BFK20 replication protein gp43. Virus Res. 163, 454–460 (2012).

29. A. Gupta, S. B. Lad, P. P. Ghodke, P. I. Pradeepkumar, K. Kondabagil, Mimivirus encodes a multifunctional primase with DNA/RNA polymerase, terminal transferase and translesion synthesis activities. Nucleic Acids Res. 47, 6932–6945 (2019).

30. S. A. MacNeill, PCNA-binding proteins in the archaea: novel functionality beyond the conserved core. Curr. Genet. 62, 527–532 (2016).

31. P. Colson, B. La Scola, A. Levasseur, G. Caetano-Anollés, D. Raoult, Mimivirus: leading the way in the discovery of giant viruses of amoebae. Nat. Rev. Microbiol. 15, 243–254 (2017).

32. A. J. Doherty, L. C. Serpell, C. P. Ponting, The Helix-Hairpin-Helix DNA-Binding Motif: A Structural Basis for Non-Sequence-Specific Recognition of DNA. Nucleic Acids Res. 24, 2488–2497 (1996).

33. L. M. Iyer, S. Balaji, E. V. Koonin, L. Aravind, Evolutionary genomics of nucleo-cytoplasmic large DNA viruses. Virus Res. 117, 156–184 (2006).

34. S. Sim, K. Hughes, X. Chen, S. L. Wolin, The Bacterial Ro60 Protein and Its Noncoding Y RNA Regulators. Annu. Rev. Microbiol. 74, 387–407 (2020).

35. C. K. Ho, L. K. Wang, C. D. Lima, S. Shuman, Structure and mechanism of RNA ligase. Structure. 12, 327–339 (2004).

36. Q. Tang, P. Wu, H. Chen, G. Li, Pleiotropic roles of the ubiquitin-proteasome system during viral propagation. Life Sci. 207, 350–354 (2018).

37. J. Murphy, J. Mahony, S. Ainsworth, A. Nauta, D. van Sinderen, Bacteriophage orphan DNA methyltransferases: insights from their bacterial origin, function, and occurrence. Appl. Environ. Microbiol. 79, 7547–7555 (2013).

38. S. Jeudy, L. Bertaux, J.-M. Alempic, A. Lartigue, M. Legendre, L. Belmudes, S. Santini, N. Philippe, L. Beucher, E. G. Biondi, S. Juul, D. J. Turner, Y. Couté, J.-M. Claverie, C. Abergel, Exploration of the propagation of transpovirons within Mimiviridae reveals a unique example of commensalism in the viral world. ISME J. 14, 727–739 (2020).

39. I. V. Agarkova, D. D. Dunigan, J. L. Van Etten, Virion-associated restriction endonucleases of chloroviruses. J. Virol. 80, 8114–8123 (2006).

40. N. Markine-Goriaynoff, L. Gillet, J. L. Van Etten, H. Korres, N. Verma, A. Vanderplasschen, Glycosyltransferases encoded by viruses. J. Gen. Virol. 85, 2741–2754 (2004).

41. F. Piacente, M. Gaglianone, M. E. Laugieri, M. G. Tonetti, The Autonomous Glycosylation of Large DNA Viruses. Int. J. Mol. Sci. 16, 29315–29328 (2015).

42. G. Hagelueken, B. R. Clarke, H. Huang, A. Tuukkanen, I. Danciu, D. I. Svergun, R. Hussain, H. Liu, C. Whitfield, J. H. Naismith, A coiled-coil domain acts as a molecular ruler to regulate O-antigen chain length in lipopolysaccharide. Nat. Struct. Mol. Biol. 22, 50–56 (2014).

43. M. Rawat, J. A. Maupin-Furlow, Redox and Thiols in Archaea. Antioxidants (Basel). 9 (2020), doi:10.3390/antiox9050381.

44. N. Joshi, F. J. Sickle, A sliding-window, adaptive, quality-based trimming tool for FastQ files (2011).

45. Y. Peng, H. C. M. Leung, S. M. Yiu, F. Y. L. Chin, IDBA-UD: a de novo assembler for single-cell and metagenomic sequencing data with highly uneven depth. Bioinformatics. 28, 1420–1428 (2012).

46. J. Alneberg, B. S. Bjarnason, I. de Bruijn, M. Schirmer, J. Quick, U. Z. Ijaz, L. Lahti, N. J. Loman, A. F. Andersson, C. Quince, Binning metagenomic contigs by coverage and composition. Nat. Methods. 11, 1144–1146 (2014).

47. D. D. Kang, F. Li, E. Kirton, A. Thomas, R. Egan, H. An, Z. Wang, MetaBAT 2: an adaptive binning algorithm for robust and efficient genome reconstruction from metagenome assemblies. PeerJ. 7, e7359 (2019).

48. C. M. K. Sieber, A. J. Probst, A. Sharrar, B. C. Thomas, M. Hess, S. G. Tringe, J. F. Banfield, Recovery of genomes from metagenomes via a dereplication, aggregation and scoring strategy. Nature Microbiology. 3 (2018), pp. 836–843.

49. D. H. Parks, M. Imelfort, C. T. Skennerton, P. Hugenholtz, G. W. Tyson, CheckM: assessing the quality of microbial genomes recovered from isolates, single cells, and metagenomes. Genome Res. 25, 1043–1055 (2015).

50. P. Jones, D. Binns, H.-Y. Chang, M. Fraser, W. Li, C. McAnulla, H. McWilliam, J. Maslen, A. Mitchell, G. Nuka, S. Pesseat, A. F. Quinn, A. Sangrador-Vegas, M. Scheremetjew, S.-Y. Yong, R. Lopez, S. Hunter, InterProScan 5: genome-scale protein function classification. Bioinformatics. 30, 1236–1240 (2014).

51. A. Biswas, R. H. J. Staals, S. E. Morales, P. C. Fineran, C. M. Brown, CRISPRDetect: A flexible algorithm to define CRISPR arrays. BMC Genomics. 17, 356 (2016).

52. L. Fu, B. Niu, Z. Zhu, S. Wu, W. Li, CD-HIT: accelerated for clustering the next-generation sequencing data. Bioinformatics. 28, 3150–3152 (2012).

53. C. Bland, T. L. Ramsey, F. Sabree, M. Lowe, K. Brown, N. C. Kyrpides, P. Hugenholtz, CRISPR recognition tool (CRT): a tool for automatic detection of clustered regularly interspaced palindromic repeats. BMC Bioinformatics. 8, 209 (2007).

54. V. A. Padilha, O. S. Alkhnbashi, S. A. Shah, A. C. P. L. F. de Carvalho, R. Backofen, CRISPRcasIdentifier: Machine learning for accurate identification and classification of CRISPR-Cas systems. Gigascience. 9 (2020), doi:10.1093/gigascience/giaa062.

55. K. Kieft, Z. Zhou, K. Anantharaman, VIBRANT: automated recovery, annotation and curation of microbial viruses, and evaluation of viral community function from genomic sequences. Microbiome. 8, 90 (2020).

56. D. Hyatt, G.-L. Chen, P. F. Locascio, M. L. Land, F. W. Larimer, L. J. Hauser, Prodigal: prokaryotic gene recognition and translation initiation site identification. BMC Bioinformatics. 11, 119 (2010).

57. T. Aramaki, R. Blanc-Mathieu, H. Endo, K. Ohkubo, M. Kanehisa, S. Goto, H. Ogata, KofamKOALA: KEGG ortholog assignment based on profile HMM and adaptive score threshold. bioRxiv (2019), p. 602110.

58. S. El-Gebali, J. Mistry, A. Bateman, S. R. Eddy, A. Luciani, S. C. Potter, M. Qureshi, L. J. Richardson, G. A. Salazar, A. Smart, E. L. L. Sonnhammer, L. Hirsh, L. Paladin, D. Piovesan, S. C. E. Tosatto, R. D. Finn, The Pfam protein families database in 2019. Nucleic Acids Res. 47, D427–D432 (2019).

59. A. L. Grazziotin, E. V. Koonin, D. M. Kristensen, Prokaryotic Virus Orthologous Groups (pVOGs): a resource for comparative genomics and protein family annotation. Nucleic Acids Res. 45, D491–D498 (2017).

60. S. R. Eddy, Accelerated Profile HMM Searches. PLoS Comput. Biol. 7, e1002195 (2011).

61. S. Nayfach, A. P. Camargo, F. Schulz, E. Eloe-Fadrosh, S. Roux, N. C. Kyrpides, CheckV assesses the quality and completeness of metagenome-assembled viral genomes. Nat. Biotechnol. 39, 578–585 (2021).

62. B. Buchfink, C. Xie, D. H. Huson, Fast and sensitive protein alignment using DIAMOND. Nat. Methods. 12, 59–60 (2014).

63. C. Camacho, G. Coulouris, V. Avagyan, N. Ma, J. Papadopoulos, K. Bealer, T. L. Madden, BLAST+: architecture and applications. BMC Bioinformatics. 10, 421 (2009).

64. D. Paez-Espino, S. Roux, I.-M. A. Chen, K. Palaniappan, A. Ratner, K. Chu, M. Huntemann, T. B. K. Reddy, J. C. Pons, M. Llabrés, E. A. Eloe-Fadrosh, N. N. Ivanova, N. C. Kyrpides, IMG/VR v.2.0: an integrated data management and analysis system for cultivated and environmental viral genomes. Nucleic Acids Res. 47, D678–D686 (2019).

65. F. Schulz, S. Roux, D. Paez-Espino, S. Jungbluth, D. A. Walsh, V. J. Denef, K. D. McMahon, K. T. Konstantinidis, E. A. Eloe-Fadrosh, N. C. Kyrpides, T. Woyke, Giant virus diversity and host interactions through global metagenomics. Nature. 578, 432–436 (2020).

66. V. A. Cantu, P. Salamon, V. Seguritan, J. Redfield, D. Salamon, R. A. Edwards, A. M. Segall, PhANNs, a fast and accurate tool and web server to classify phage structural proteins. bioRxiv (2020), p. 2020.04.03.023523.

67. L. Zimmermann, A. Stephens, S.-Z. Nam, D. Rau, J. Kübler, M. Lozajic, F. Gabler, J. Söding, A. N. Lupas, V. Alva, A Completely Reimplemented MPI Bioinformatics Toolkit with a New HHpred Server at its Core. J. Mol. Biol. 430, 2237–2243 (2018).

68. J. R. Grant, P. Stothard, The CGView Server: a comparative genomics tool for circular genomes. Nucleic Acids Res. 36, W181–W184 (2008).

69. H. Bin Jang, B. Bolduc, O. Zablocki, J. H. Kuhn, S. Roux, E. M. Adriaenssens, J. R. Brister, A. M. Kropinski, M. Krupovic, R. Lavigne, D. Turner, M. B. Sullivan, Taxonomic assignment of uncultivated prokaryotic virus genomes is enabled by gene-sharing networks. Nat. Biotechnol. 37, 632–639 (2019).

70. T. Nepusz, H. Yu, A. Paccanaro, Detecting overlapping protein complexes in protein-protein interaction networks. Nat. Methods. 9, 471–472 (2012).

71. A. J. Enright, S. Van Dongen, C. A. Ouzounis, An efficient algorithm for large-scale detection of protein families. Nucleic Acids Res. 30, 1575–1584 (2002).

72. E. W. Sayers, T. Barrett, D. A. Benson, S. H. Bryant, K. Canese, V. Chetvernin, D. M. Church, M. DiCuccio, R. Edgar, S. Federhen, M. Feolo, L. Y. Geer, W. Helmberg, Y. Kapustin, D. Landsman, D. J. Lipman, T. L. Madden, D. R. Maglott, V. Miller, I. Mizrachi, J. Ostell, K. D. Pruitt, G. D. Schuler, E. Sequeira, S. T. Sherry, M. Shumway, K. Sirotkin, A. Souvorov, G. Starchenko, T. A. Tatusova, L. Wagner, E. Yaschenko, J. Ye, Database resources of the National Center for Biotechnology Information. Nucleic Acids Res. 37, D5–15 (2009).

73. B. Al-Shayeb, R. Sachdeva, L.-X. Chen, F. Ward, P. Munk, A. Devoto, C. J. Castelle, M. R. Olm, K. Bouma-Gregson, Y. Amano, C. He, R. Méheust, B. Brooks, A. Thomas, A. Lavy, P. Matheus-Carnevali, C. Sun, D. S. A. Goltsman, M. A. Borton, A. Sharrar, A. L. Jaffe, T. C. Nelson, R. Kantor, R. Keren, K. R. Lane, I. F. Farag, S. Lei, K. Finstad, R. Amundson, K. Anantharaman, J. Zhou, A. J. Probst, M. E. Power, S. G. Tringe, W.-J. Li, K. Wrighton, S. Harrison, M. Morowitz, D. A. Relman, J. A. Doudna, A.-C. Lehours, L. Warren, J. H. D. Cate, J. M. Santini, J. F. Banfield, Clades of huge phages from across Earth’s ecosystems. Nature. 578, 425–431 (2020).

74. P. Shannon, A. Markiel, O. Ozier, N. S. Baliga, J. T. Wang, D. Ramage, N. Amin, B. Schwikowski, T. Ideker, Cytoscape: a software environment for integrated models of biomolecular interaction networks. Genome Res. 13, 2498–2504 (2003).

75. RStudio Team, RStudio: Integrated Development Environment for R (2019), (available at http://www.rstudio.com/).

76. R Core Team, R: A Language and Environment for Statistical Computing (2020), (available at https://www.R-project.org/).

77. B. Rudis, D. Gandy, waffle: Create waffle chart visualizations in R (2016), (available at https://CRAN.R-project.org/package=waffle).

78. V. De Anda, I. Zapata-Peñasco, J. Blaz, A. C. Poot-Hernández, B. Contreras-Moreira, M. González-Laffitte, N. Gámez-Tamariz, M. Hernández-Rosales, L. E. Eguiarte, V. Souza, Understanding the Mechanisms Behind the Response to Environmental Perturbation in Microbial Mats: A Metagenomic-Network Based Approach. Front. Microbiol. 9, 2606 (2018).

79. J. Guglielmini, A. C. Woo, M. Krupovic, P. Forterre, M. Gaia, Diversification of giant and large eukaryotic dsDNA viruses predated the origin of modern eukaryotes. Proc. Natl. Acad. Sci. U. S. A. 116, 19585–19592 (2019).

80. K. Katoh, D. M. Standley, MAFFT multiple sequence alignment software version 7: improvements in performance and usability. Mol. Biol. Evol. 30, 772–780 (2013).

81. B. Q. Minh, H. A. Schmidt, O. Chernomor, D. Schrempf, M. D. Woodhams, A. von Haeseler, R. Lanfear, IQ-TREE 2: New Models and Efficient Methods for Phylogenetic Inference in the Genomic Era. Mol. Biol. Evol. 37, 1530–1534 (2020).

82. S. Kalyaanamoorthy, B. Q. Minh, T. K. F. Wong, A. von Haeseler, L. S. Jermiin, ModelFinder: fast model selection for accurate phylogenetic estimates. Nat. Methods. 14, 587–589 (2017).

83. I. Letunic, P. Bork, Interactive Tree Of Life (iTOL) v5: an online tool for phylogenetic tree display and annotation. Nucleic Acids Res. 49, W293–W296 (2021).

