## Supplementary Figures for "Unique viruses that infect Archaea related to eukaryotes"

**Supplementary Information for Rambo, De Anda, Langwig and Baker 2021, "Unique viruses that infect Archaea related to eukaryotes".**

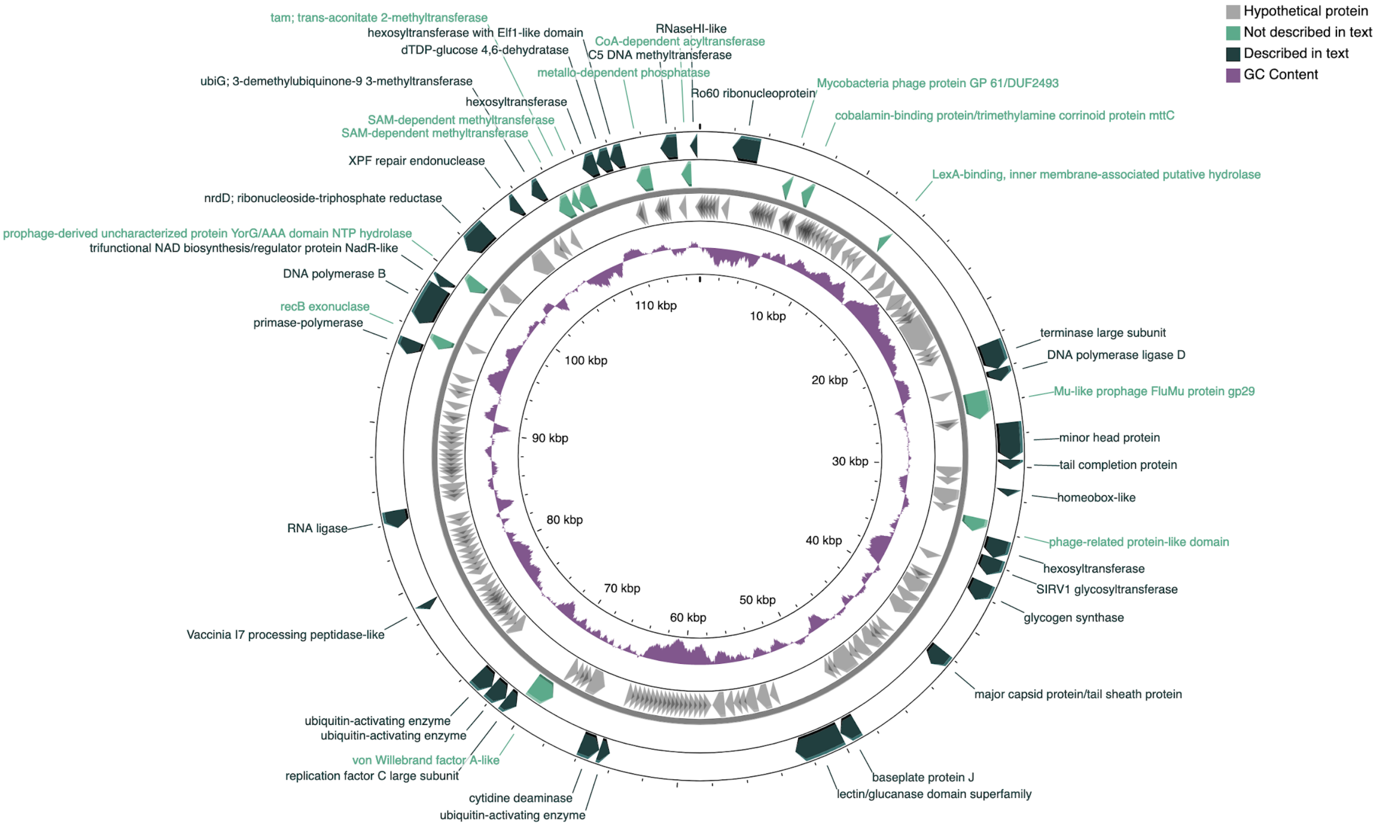


**Fig. S1. Genomic architecture of the complete Helarchaeota virus Nidhogg Meg22_1012.** From outside to center: genes described in the main text, genes with homologs not described in the main text, hypothetical proteins, GC content, genome size ruler. Arrows pointing left indicate (-) sense, while those pointing right indicate (+) sense.

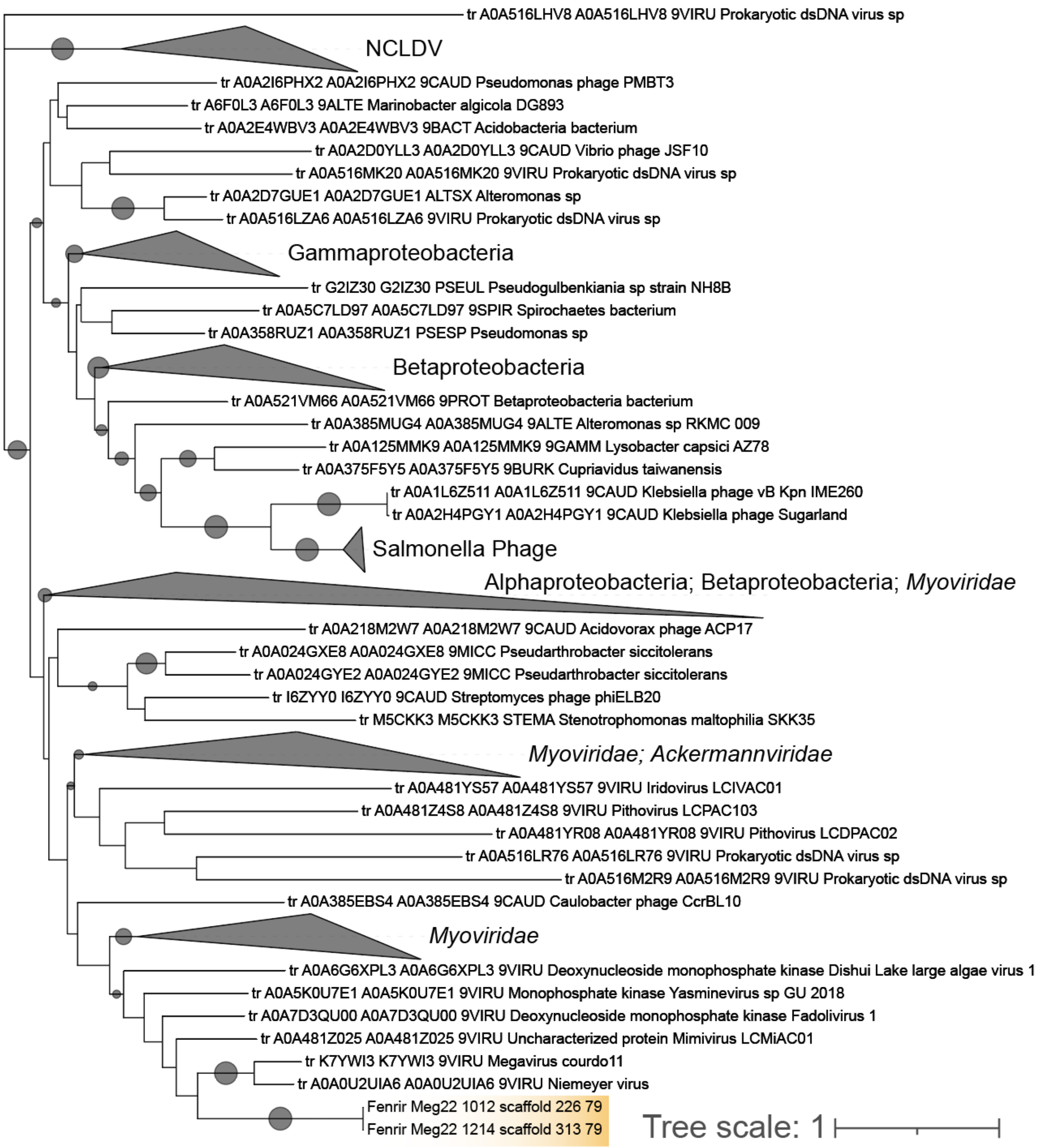


**Fig. S2. Deoxynucleotide monophosphate kinase phylogeny.** A phylogenetic tree of 241 deoxynucleotide/side monophosphate kinase sequences from viruses and bacteria. Circles on branches indicate BOOSTER supports >= 70%. Lokiarchaeota virus Fenrir Meg22_1012 and Meg22_1214 sequences are highlighted in gold. The phylogeny was inferred using the LG model with fixed base frequencies and 1,000 rapid bootstraps.

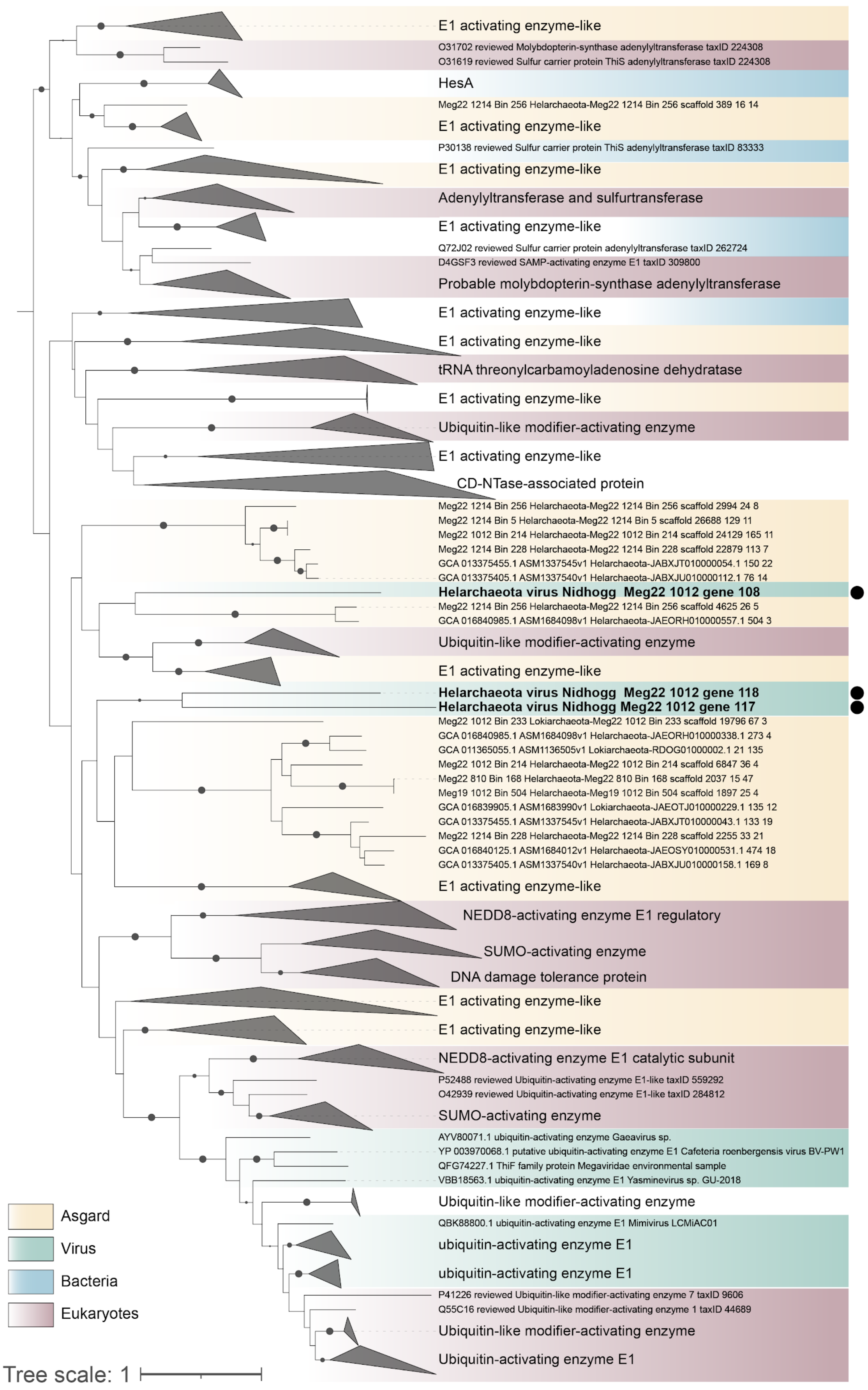
E1 activating enzyme-like
O31702 reviewed Molybdopterin-synthase adenylyltransferase taxID 224308
O31619 reviewed Sulfur carrier protein ThiS adenylyltransferase taxID 224308
HesA
Meg22 1214 Bin 256 Helarchaeota-Meg22 1214 Bin 256 scaffold 389 16 14
E1 activating enzyme-like
P30138 reviewed Sulfur carrier protein ThiS adenylyltransferase taxID 83333
E1 activating enzyme-like
Adenylyltransferase and sulfurtransferase
E1 activating enzyme-like
Q72J02 reviewed Sulfur carrier protein adenylyltransferase taxID 262724
D4GSF3 reviewed SAMP-activating enzyme E1 taxID 309800
Probable molybdopterin-synthase adenylyltransferase
E1 activating enzyme-like
E1 activating enzyme-like
tRNA threonylcarbamoyladenosine dehydratase
E1 activating enzyme-like
Ubiquitin-like modifier-activating enzyme
E1 activating enzyme-like
CD-NTase-associated protein
Meg22 1214 Bin 256 Helarchaeota-Meg22 1214 Bin 256 scaffold 2994 24 8
Meg22 1214 Bin 5 Helarchaeota-Meg22 1214 Bin 5 scaffold 26688 129 11
Meg22 1012 Bin 214 Helarchaeota-Meg22 1012 Bin 214 scaffold 24129 165 11
Meg22 1214 Bin 228 Helarchaeota-Meg22 1214 Bin 228 scaffold 22879 113 7
GCA 013375455.1 ASM1337545v1 Helarchaeota-JABXJT010000054.1 150 22
GCA 013375405.1 ASM1337540v1 Helarchaeota-JABXJU010000112.1 76 14
Helarchaeota virus Nidhogg Meg22 1012 gene 108
Meg22 1214 Bin 256 Helarchaeota-Meg22 1214 Bin 256 scaffold 4625 26 5
GCA 016840985.1 ASM1684098v1 Helarchaeota-JAEORH010000557.1 504 3
Ubiquitin-like modifier-activating enzyme
E1 activating enzyme-like
Helarchaeota virus Nidhogg Meg22 1012 gene 118
Helarchaeota virus Nidhogg Meg22 1012 gene 117
Meg22 1012 Bin 233 Lokiarchaeota-Meg22 1012 Bin 233 scaffold 19796 67 3
GCA 016840985.1 ASM1684098v1 Helarchaeota-JAEORH010000338.1 273 4
GCA 011365055.1 ASM1136505v1 Lokiarchaeota-RDOG01000002.1 21 135
Meg22 1012 Bin 214 Helarchaeota-Meg22 1012 Bin 214 scaffold 6847 36 4
Meg22 810 Bin 168 Helarchaeota-Meg22 810 Bin 168 scaffold 2037 15 47
Meg19 1012 Bin 504 Helarchaeota-Meg19 1012 Bin 504 scaffold 1897 25 4
GCA 016839905.1 ASM1683990v1 Lokiarchaeota-JAEOTJ010000229.1 135 12
GCA 013375455.1 ASM1337545v1 Helarchaeota-JABXJT010000043.1 133 19
Meg22 1214 Bin 228 Helarchaeota-Meg22 1214 Bin 228 scaffold 2255 33 21
GCA 016840125.1 ASM1684012v1 Helarchaeota-JAEOSY010000531.1 474 18
GCA 013375405.1 ASM1337540v1 Helarchaeota-JABXJU010000158.1 169 8
E1 activating enzyme-like
NEDD8-activating enzyme E1 regulatory
SUMO-activating enzyme
DNA damage tolerance protein
E1 activating enzyme-like
E1 activating enzyme-like
NEDD8-activating enzyme E1 catalytic subunit
P52488 reviewed Ubiquitin-activating enzyme E1-like taxID 559292
O42939 reviewed Ubiquitin-activating enzyme E1-like taxID 284812
SUMO-activating enzyme
AYV80071.1 ubiquitin-activating enzyme Gaeavirus sp.
YP 003970068.1 putative ubiquitin-activating enzyme E1 Cafeteria roenbergensis virus BV-PW1
QFG74227.1 ThiF family protein Megaviridae environmental sample
VBB18563.1 ubiquitin-activating enzyme E1 Yasminevirus sp. GU-2018
Ubiquitin-like modifier-activating enzyme
QBK88800.1 ubiquitin-activating enzyme E1 Mimivirus LCMiAC01
ubiquitin-activating enzyme E1
ubiquitin-activating enzyme E1
P41226 reviewed Ubiquitin-like modifier-activating enzyme 7 taxID 9606
Q55C16 reviewed Ubiquitin-like modifier-activating enzyme 1 taxID 44689
Ubiquitin-like modifier-activating enzyme
Ubiquitin-activating enzyme E1
Asgard
Virus
Bacteria
Eukaryotes
Tree scale: 1

**Fig. S3. Ubiquitin-activating enzyme phylogeny.** A phylogenetic tree of 368 ubiquitin-activating enzyme (E1) protein sequences from archaea, bacteria, eukaryotes, and viruses (taxa are labeled with background colors). Three E1-like protein sequences were identified in Nidhogg viruses and these are labeled with black circles and bold text. Arched lines show the connections between Nidhogg virus sequences and their Helarchaeota host. This phylogeny was inferred using the LG+R8 model with 1,000 ultrafast bootstraps and optimization by nearest neighbor interchange (-bb 1000 -bnni). The tree is comprised of protein sequences belonging to the NEDD8-activating enzyme E1 catalytic subunit family (n = 11, IPR030468), ubiquitin-activating E1 enzyme (n = 218, IPR035985), viral sequences obtained from NCBI (n = 14), and sequences derived from Lokiarchaeota and Helarchaeota (n = 125).
